# Segregation and integration between visuo-spatial attention and semantic memory across large-scale brain networks

**DOI:** 10.64898/2026.07.21.739848

**Authors:** Michelangelo Tani, Sandeep Kaur, Maria Ciociola, Maria Bianca Muneghina, Alma Cecconi, Valentina Sulpizio, Antonello Baldassarre, Paolo Capotosto, Gaspare Galati

## Abstract

Visuo-spatial attention and semantic memory are supported by distinct, largely competing brain networks, yet they are frequently engaged together in daily life. How these networks interact during combined tasks remains unexplored. We conducted a factorial fMRI experiment in which 25 participants performed a novel task requiring visuo-spatial attention, semantic judgment, or both. Covert shifts of attention towards cued lateral location activated bilateral parietal (PEF), frontal eye fields (FEF), and anterior insula. Categorizing a word as referring to a living or non-living entity activated a lateral parietal region (LaP) between the dorsal tip of the angular gyrus and the lateral bank of the intraparietal sulcus, along with the left inferior frontal gyrus (IFG), superior temporal sulcus, inferior temporal lobe and bilateral anterior insula. Notably, the anterior insula was active in both conditions. Dynamic causal modeling showed excitatory-inhibitory parieto-frontal loops driving attention (PEF→FEF) and semantic processing (LaP→IFG) separately, but in the combined condition frontal-to-parietal feedback became excitatory, reflecting stronger integration, with the anterior insula driving overall connectivity. These findings identify LaP as a novel region supporting semantic processing of linguistic stimuli, and highlight the anterior insula as a key hub integrating attentional and semantic networks under concurrent cognitive demands.

**Highlights:** Visuo-spatial attention and semantic memory rely on segregated parieto-frontal circuits. However, concurrent demands induce a large-scale reconfiguration centered on the left anterior insula, which acts as an integrative hub between the two networks.

## 1. Introduction

Visuo-spatial attention and semantic memory are among the most extensively studied cognitive functions in human neuroscience. A large body of evidence indicates that visuo-spatial orienting relies on a distributed frontoparietal system, commonly referred to as the dorsal attention network (DAN), whose core nodes include the intraparietal sulcus (IPS) and the frontal eye fields (FEF) (Corbetta & Shulman, 2002). In contrast, semantic processing is supported by a large-scale widely distributed network that is deeply intertwined with the default mode network (DMN) (Nieberlein et al., 2024; Thye et al., 2025). This system has also been characterized as encompassing a semantic control network involving lateral prefrontal and posterior temporal regions (Noonan et al., 2013; Ralph et al., 2017; Herault et al., 2024; Dumitrescu et al., 2025), with relevant sensory modality, attribute-specific and task-related variability (Kiefer & Pulvermüller, 2012; Thompson-Schill et al., 2003; Binder et al., 2009; Jedidi et al., 2021; Blackett et al., 2022; Hodgson et al., 2024; Quartetti et al., 2025; Radman et al., 2026).

From a network perspective, attentional and semantic processes have often been associated with partially segregated, and in some circumstances competing, large-scale systems. Resting-state studies have consistently shown anticorrelated activity patterns between the DAN and the DMN (Fox et al., 2005). In contrast, task-based studies have reported preferential recruitment of these networks during externally directed visuo-spatial processing and internally oriented memory-related operations.

However, in everyday behavior, attention and memory rarely operate in isolation: reading, visual search, object recognition, and social interaction typically require the simultaneous allocation of spatial attention and access to semantic knowledge. Understanding how these networks interact under concurrent demands is therefore a fundamental question for systems neuroscience. While the neural substrates of each process have been extensively characterized separately, considerably less is known about the mechanisms enabling their integration — for instance, whether their interaction is coordinated by a third source such as the cingulo-opercular network (Dosenbach et al., 2007; Power et al., 2011).

Previous work from our group provided causal evidence for a functional dissociation between parietal regions involved in attention and semantic processing. Using a combined transcranial magnetic stimulation (TMS) and electroencephalography (EEG) approach, we showed that behavioral performance and several local and global EEG markers (e.g., the parietal event-related de-synchronization of the alpha rhythms and the topography of the EEG microstates) were selectively disrupted when stimulating the intraparietal sulcus during a visuo-spatial attentional task, and when stimulating the angular gyrus during a semantic memory retrieval task (Capotosto et al., 2017; Croce et al., 2018). More recently, we employed a task requiring the simultaneous engagement of both processes and found that inhibition of either region impaired behavioral performance (Capotosto et al., 2026). These findings indicate that attention and semantic systems can be experimentally dissociated and contribute to the successful execution of combined demands, yet must interact when concurrently required.

The present study aimed to characterize the large-scale neural architecture supporting visuo-spatial attention and semantic memory, both in isolation and under concurrent demands. To this end, we developed a factorial fMRI paradigm in which participants performed tasks involving visuo-spatial attention, semantic categorization, or a combination of both. First, we sought to identify the cortical regions selectively associated with each function. Second, we investigated whether the concurrent engagement of attention and semantic memory recruits additional neural mechanisms beyond those observed during isolated processing. Finally, we used Dynamic Causal Modeling (DCM) to examine how effective connectivity is reconfigured when attentional and semantic demands must be integrated within a single behavioral context.

## 2. Methods

### 2.1 Participants

Twenty-five right-handed volunteers participated in the study (15 females; age: 24.5 ± 2.5 years old). Participants were native Italian speakers with normal or corrected-to-normal vision and no neurological or psychiatric disorder. Sample size was based on established guidelines indicating 20–25 participants provide adequate power for medium-to-large effects in novel fMRI paradigms (Desmond & Glover, 2002; Murphy & Garavan, 2004). The study received ethical approval from the Ethics Committee of the Regione Lazio Area 5, Italy (approval 0003016.28-02-2024). All participants gave written informed consent and volunteered without compensation.

### 2.2 Experimental design

We employed a 2×2 factorial design with two factors — visuo-spatial attention and semantic memory, either present or absent — yielding four conditions. In each trial (Fig. 1), participants were shown a target letter sequence preceded by a central cue and performed a discrimination task. In the semantic conditions, the letter sequence was an Italian word, and participants performed a living/non-living categorization; in the non-semantic conditions, the letter sequence was a non-word, and participants judged whether it contained the letter “A”. In the attentional conditions, the target letter sequence appeared either to the left or to the right of the central fixation point, and a central cue informed about the side in which the target would appear, with an 80% validity (Posner, 1980); in the non-attentional conditions, the target letter appeared centrally, and the cue was non-informative.

**Figure 1.**
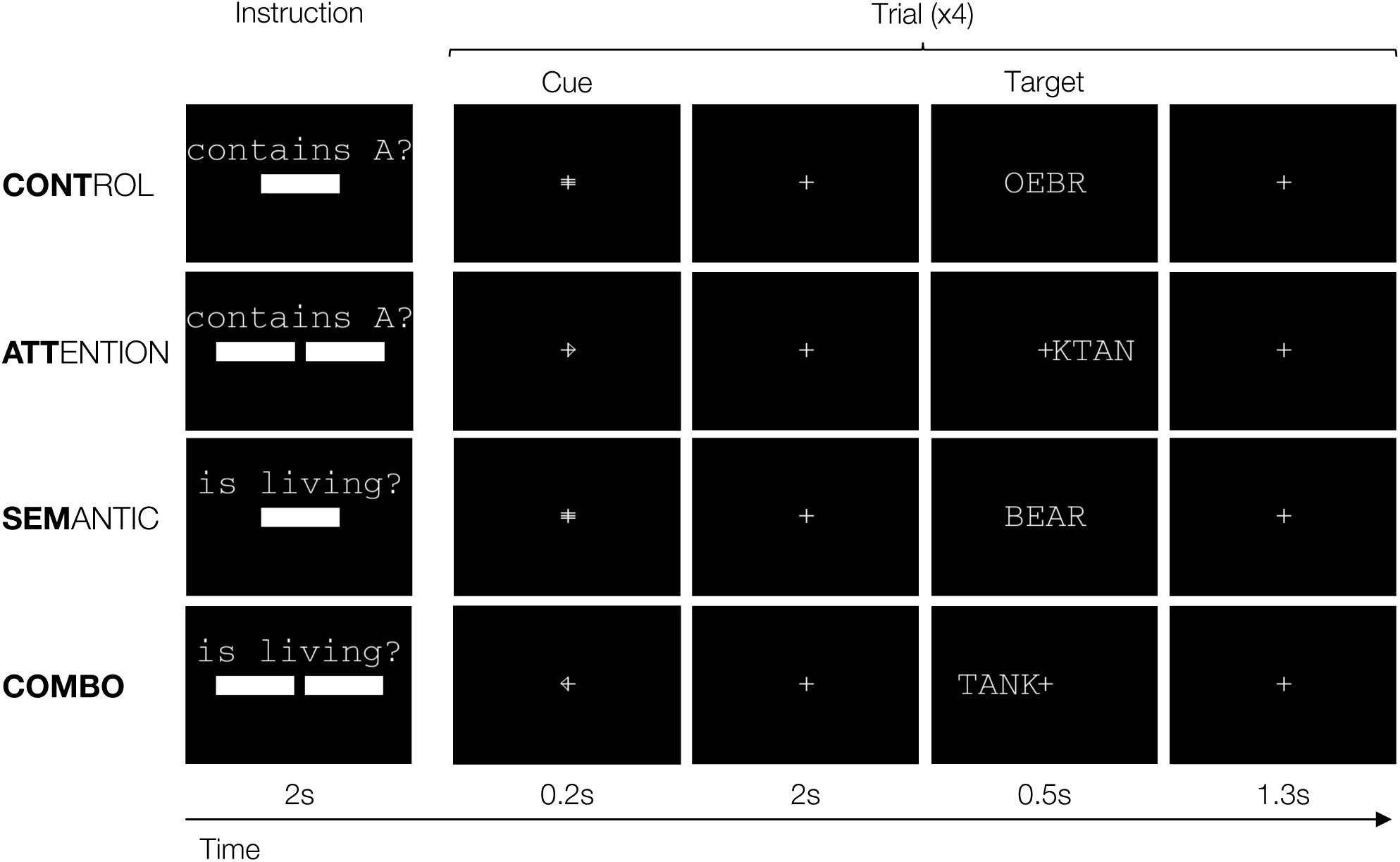
Experimental paradigm. The experiment employed a 2×2 factorial block design crossing two factors — visuo-spatial attention (present or absent) and semantic processing (present or absent) — yielding four conditions: CONT (neither factor), ATT (attention only), SEM (semantic only), and COMBO (both factors). Each row illustrates one condition. When the semantic factor was present, participants judged whether a word referred to a living or non-living entity (SEM, COMBO); when absent, they judged whether a non-word contained the letter A (CONT, ATT). When the visuo-spatial attention factor was present, targets appeared laterally with a directional cue valid on 80% of trials (ATT, COMBO); when absent, targets appeared centrally preceded by a neutral cue (CONT, SEM). Each block comprised 4 trials and was preceded by a 2-s instruction period consisting of 1 s of written task instruction with one or two white rectangles indicating the possible target location, followed by 1 s of fixation. Each trial began with a 200 ms directional cue superimposed on the fixation cross — a neutral symbol (=) when targets would appear centrally or an arrow (< or >) when targets would appear laterally — followed by a 2 s cue-target interval, a 500 ms target stimulus, and a 1300 ms response window. Participants responded by pressing a “yes” button with the right index finger or a “no” button with the right middle finger. Stimuli are enlarged for illustrative purposes; target stimuli were presented in Italian.

The four conditions were:

1. Attention-only (ATT): covert attention to the cued hemifield, discriminating a laterally presented non-word for the letter “A”;
2. Semantic-only (SEM): discriminating whether a centrally presented word referred to a living or non-living entity;
3. Combined (COMBO): covert attention to the cued hemifield, discriminating whether a laterally presented word referred to a living or non-living entity;
4. Control (CONT): discriminating whether a centrally presented non-word contained the letter “A”.

### 2.3 Stimuli and task

Target stimuli comprised 400 items: 200 four-letter Italian words (100 living, 100 non-living) and 200 four-letter non-words (100 containing “A”, 100 not). Words were drawn from the CoLFIS frequency lexicon (Bertinetto & Loporcaro, 2005), the same set used by Capotosto et al. (2023). Non-words were created by rearranging word letters into meaningless combinations (e.g., “ROSA” → “SROA”) to avoid semantic elaboration; if a source word contained two “A”s, one was replaced with a different vowel. Stimuli were presented in uppercase Courier font (0.3° height, 1.2° width) on a black background.

Trials were arranged into a block design. Each block included 4 trials and was introduced by a 2s instruction period, comprising 1s of written task instruction (“Is it a living entity?” for SEM and COMBO blocks; “Does it contain the letter A?” for ATT and CONT blocks), followed by 1s of fixation. Instruction included a single central rectangle (in SEM and CONT blocks) or two lateral rectangles (in COMBO and ATT blocks), indicating possible target locations.

Participants fixated a central cross (0.3°) throughout the task. Each trial began with a 200 ms cue indicating the target location: a leftward arrow (<), a rightward arrow (>), or a neutral symbol (=). Arrows subtended 0.15° x 0.3° of visual angle and were presented laterally, with their open edges superimposed on the vertical segment of the fixation cross. After a 2000ms cue-target interval, a word/non-word appeared for 500ms, followed by a 1300ms response window. In ATT and COMBO blocks, stimuli appeared within the parafovea (eccentricity: from ±0.15° to ±1.35° horizontally from fixation), with 80% valid and 20% invalid arrows cueing. In SEM and CONT blocks, stimuli appeared centrally with a neutral cue. Participants responded via button box, pressing “Yes” with the index finger or “No” with the middle finger of the right hand.

### 2.4 Apparatus

Data were acquired on a Magnetom Prisma 3T scanner (Siemens, Erlangen, Germany) at the Neuroimaging Laboratory, Santa Lucia Foundation, Rome, with a 32-channel head coil. Stimuli were generated in PsychoPy 2024.1.5 (Open Science Tools Ltd, University of Nottingham) on a PC outside the scanner room and projected via an NEC PA572W LCD projector (1280×800, 60Hz) onto a screen behind the bore (33.6°×19.5° visual angle; distance = 77cm), viewed via a mirror on the head coil. Timing was locked to the scanner trigger via an optical line (FIU-932, Current Designs, Philadelphia, USA). Stimulus timing was controlled at the refresh rate (60 Hz) and responses were recorded via a two-button fiber-optic pad (HHSC-2×4-CR, Current Designs), logged by PsychoPy for reaction time and accuracy at sub-millisecond resolution. Training was administered prior to the scanning session on a MacBook Air (M3) running PsychoPy 2024.1.5.

### 2.5 Procedure

Participants completed the Edinburgh Handedness Inventory (Oldfield, 1971) and an MRI eligibility health screening, then received informed consent and task instructions, followed by a training run to familiarized with the task. Participants then lay supine with the head coil and mirror mounted, comfort padding applied to minimize head movement, and a cushion was placed under the legs for comfort and the response box under the right hand.

For each participant, a pseudorandom, optimally counterbalanced trial list was generated across 20 randomization iterations, comprising five runs each containing all 400 stimuli (each presented once). Each run had 80 trials (40 words, 40 non-words), conditions counterbalanced within runs; one run was used for training, four for the experimental session.

Each of the four experimental runs comprised 20 blocks in five sets, with the four conditions pseudorandomly ordered within each set. Across the 25 sets in four runs, 24 possible condition-order permutations were used, one repeated. A 16s rest period preceded and followed each set to estimate baseline. The task lasted ∼28 minutes; the full session, including instructions, training, and preparation, lasted ∼70 minutes.

### 2.6 Image acquisition

Structural and functional images were acquired with the following imaging protocols:

1. T1-weighted MPRAGE: 176 slices, 1mm³ isotropic resolution, FOV 256×256mm, TR=2500ms, TE=1.67ms, inversion time=1080ms, flip angle=8°.
2. T2-weighted 3D SPACE: variable flip angle refocusing (nominal 120°), 176 slices, 1mm³ isotropic resolution, FOV 256×240mm, TR=3200ms, TE=564ms.
3. Five functional whole-brain T2*-weighted gradient-echo EPI runs: multiband factor 6, 2.4mm³ isotropic voxel size, 60 slices, FOV 88×88mm², TR=800ms, TE=30ms, flip angle=52°, no in-plane acceleration (Xu et al., 2013; Feinberg et al., 2010; Moeller et al., 2010).
4. Two spin-echo EPI field-mapping volumes with opposite phase encoding, no multiband acceleration, matching geometry and sampling properties of functional scans (TE=80ms, TR=7060ms, flip angle=90°).

Both structural scans used perspective motion correction and selective reacquisition via interleaved 3D EPI navigators (Tisdall et al., 2012; Hess et al., 2011). Participants underwent four task runs (583 volumes each) and one resting-state run (450 volumes), not analyzed here.

### 2.7 Image preprocessing

Results reported in this manuscript were obtained from preprocessing using fMRIPrep 25.0.0 (Esteban et al., 2019; Esteban et al., 2018; RRID: SCR_016216), which is based on Nipype 1.9.2 (Gorgolewski et al., 2011; Gorgolewski et al., 2018; RRID: SCR_002502).

#### 2.7.1 Anatomical data preprocessing

The T1w image was corrected for intensity non-uniformity (INU) with N4BiasFieldCorrection (Tustison et al., 2010), distributed with ANTs 2.5.4 (Avants et al., 2008, RRID: SCR_004757), and used as T1w-reference throughout the workflow. The T1w-reference was then skull-stripped using a Nipype implementation of the antsBrainExtraction.sh workflow (from ANTs), with OASIS30ANTs as the target template. Brain tissue segmentation of cerebrospinal fluid (CSF), white-matter (WM), and gray-matter (GM) was performed on the brain-extracted T1w using fast (FSL (version unknown), RRID: SCR_002823, Zhang, Brady, and Smith 2001). Brain surfaces were reconstructed using recon-all (FreeSurfer 7.3.2, RRID: SCR_001847, Dale, Fischl, and Sereno 1999), and the brain mask estimated previously was refined with a custom variation of the method to reconcile ANTs-derived and FreeSurfer-derived segmentations of the cortical gray matter of Mindboggle (RRID: SCR_002438, Klein et al. 2017). A T2-weighted image was used to improve pial surface refinement. Brain surfaces were reconstructed using recon-all (FreeSurfer 7.3.2, RRID: SCR_001847, Dale, Fischl, and Sereno 1999), and the brain mask estimated previously was refined with a custom variation of the method to reconcile ANTs-derived and FreeSurfer-derived segmentations of the cortical gray matter of Mindboggle (RRID: SCR_002438, Klein et al. 2017). Volume-based spatial normalization to two standard spaces (MNI152NLin6Asym, MNI152NLin2009cAsym) was performed through nonlinear registration with antsRegistration (ANTs 2.5.4), using brain-extracted versions of both T1w reference and the T1w template. The following templates were selected for spatial normalization and accessed with TemplateFlow (24.2.2, Ciric et al. 2022): FSL’s MNI ICBM 152 non-linear 6th Generation Asymmetric Average Brain Stereotaxic Registration Model [Evans et al. (2012), RRID: SCR_002823; TemplateFlow ID: MNI152NLin6Asym], ICBM 152 Nonlinear Asymmetrical template version 2009c [Fonov et al. (2009), RRID: SCR_008796; TemplateFlow ID: MNI152NLin2009cAsym]. Grayordinate “dscalar” files containing 91k samples were resampled onto fsLR using the Connectome Workbench (Glasser et al., 2013).

#### 2.7.2 Functional data preprocessing

For each of the 5 BOLD runs acquired per subject (across all tasks and sessions), the following preprocessing was performed. First, a reference volume was generated using a custom fMRIPrep methodology for head motion correction. Head-motion parameters with respect to the BOLD reference (transformation matrices and six corresponding rotation and translation parameters) are estimated before any spatiotemporal filtering using mcflirt (FSL, Jenkinson et al. 2002). The estimated fieldmap was then rigidly registered to the target EPI (echo-planar imaging) reference run. The field coefficients were mapped onto the reference EPI using the transform. The BOLD reference was then co-registered to the T1w reference using bbregister (FreeSurfer), which implements boundary-based registration (Greve & Fischl, 2009). Co-registration was configured with six degrees of freedom. The aligned T2w image was used for initial co-registration.

Several confounding time series were calculated from the preprocessed BOLD: framewise displacement (FD), DVARS, and three region-wise global signals. FD and DVARS were calculated for each functional run using Nipype implementations (following the definitions by Power et al., 2014). The three global signals are extracted within the CSF, the WM, and the whole-brain masks. Additionally, a set of physiological regressors was extracted to enable component-based noise correction (CompCor; Behzadi et al., 2007). Principal components are estimated after high-pass filtering the preprocessed BOLD time series (using a discrete cosine filter with a 128s cut-off) for the two CompCor variants: temporal (tCompCor) and anatomical (aCompCor). tCompCor components are then calculated from the top 2% variable voxels within the brain mask. For aCompCor, three probabilistic masks (CSF, WM, and combined CSF+WM) are generated in anatomical space. The implementation differs from that of Behzadi et al. (2007) in that, instead of eroding the masks by 2 pixels in BOLD space, a mask of pixels likely containing a GM volume fraction is subtracted from the aCompCor masks. This mask is obtained by dilating a GM mask extracted from FreeSurfer’s aseg segmentation, ensuring components are not extracted from voxels containing a minimal fraction of GM. Finally, these masks are resampled into BOLD space and binarized using a threshold of 0.99 (as in the original implementation). Components are also calculated separately within the WM and CSF masks. For each CompCor decomposition, the k components with the largest singular values are retained, such that the retained components’ time series are sufficient to explain 50 percent of variance across the nuisance mask (CSF, WM, combined, or temporal). The remaining components are dropped from consideration. The head-motion estimates calculated in the correction step were also placed within the corresponding confounds file. The confound time series derived from head motion estimates and global signals were expanded by including temporal derivatives and quadratic terms for each (Satterthwaite et al., 2013). Frames that exceeded a threshold of 0.5 mm FD or 1.5 standardized DVARS were annotated as motion outliers. Additional nuisance time series are calculated by principal components analysis of the signal within a thin band (crown) of voxels around the edge of the brain, as proposed by Patriat, Reynolds, and Birn (2017).

The BOLD time-series were resampled onto the left/right-symmetric template “fsLR” using the Connectome Workbench (Glasser et al., 2013). A “goodvoxels” mask was applied during volume-to-surface sampling in fsLR space, excluding voxels whose time series have a locally high coefficient of variation. Grayordinates files (Glasser et al., 2013) containing 91k samples were also generated, with surface data transformed directly to the fsLR space and subcortical data transformed to 2 mm resolution MNI152NLin6Asym space. All resamplings can be performed with a single interpolation step by composing all the pertinent transformations (i.e., head-motion transform matrices, susceptibility distortion correction when available, and co-registrations to anatomical and output spaces). Gridded (volumetric) resamplings were performed using nitransforms, configured with cubic B-spline interpolation. Many internal operations of fMRIPrep use Nilearn 0.11.1 (Abraham et al., 2014, RRID: SCR_001362), primarily in the functional processing workflow. For more details on the pipeline, see the workflows section of fMRIPrep’s documentation.

### 2.8 Image analysis

Functional images were analyzed in surface-based fsLR space, vertex-by-vertex, using a general linear model (GLM) in SPM12 (Wellcome Center for Human Neuroimaging). Time series were spatially smoothed (6mm FWHM Gaussian kernel) before GLM estimation, balancing signal-to-noise ratio with surface-data suitability. Each block was modeled as a boxcar function (onset at block-instruction start, offset at the next block’s instruction onset, ∼18s) convolved with the canonical HRF, with separate regressors for ATT, SEM, COMBO, and CONT. Six nuisance regressors were included: framewise displacement, the first five anatomical CompCor components, and a constant term.

First-level parameter estimates yielded subject-level contrasts for the main effect of attention [(ATT+COMBO)-(SEM+CONT)], the main effect of semantic memory [(SEM+COMBO)-(ATT+CONT)], and their interaction [(COMBO+CONT)-(ATT+SEM)] and its reverse. These were entered into second-level one-sample t-tests for group statistical maps, masked for vertices with above-baseline average activity and thresholded at *p*<0.05 cluster-corrected via topological false discovery rate (Chumbley et al., 2010), with a cluster-forming threshold of *p*<0.001 uncorrected at the vertex level.

### 2.9 Dynamic Causal Modeling

Dynamic causal modeling (DCM; Buxton et al., 1998; Friston et al., 2000) examined effective-connectivity modulations induced by the combined task within and across the networks involved in visuospatial attention and semantic memory. DCM uses an extended balloon model linking hidden neuronal dynamics to BOLD signal, with a bilinear approximation expressing neural state changes as functions of endogenous connectivity (A matrix), condition-driven modulation of connections (B matrix), and direct driving inputs (C matrix).

Connectivity nodes were derived from group-level statistical maps of the main effects of attention and semantic memory, based on anatomical location and consistency with prior literature; given the linguistic stimuli, we focused on left-hemisphere intrahemispheric connectivity. The A matrix, encoding baseline connectivity estimated from inter-block rest periods, was modeled using structural connectivity considerations combining proximity and known long-range fasciculi (see Results for node and connection details).

Where possible, nodes were defined individually by selecting participant-level activation clusters (main effects, reduced 4mm FWHM smoothing); watershed segmentation (Mangan & Whitaker, 1999) delineated regions when clusters contained multiple local maxima. When a participant lacked significant activation in a node, the group-level region was used, preserving network coverage while retaining individual variability where possible.

For each node, a representative time series was extracted from unsmoothed preprocessed data as the first eigenvariate after regressing out effects of no interest; CONT epochs were treated as effects of no interest.

Extracted time series underwent first-level DCM estimation, with the model inverted per participant using the Variational Laplace scheme (Friston et al., 2007), maximizing variational free energy as a model-evidence proxy. Models explaining less than 10% of BOLD variance were flagged as poorly inverted (Zeidman et al., 2019a).

Group-level inference used Parametric Empirical Bayes (PEB; Friston et al., 2015, 2016), pooling individual DCMs into a second-level model accounting for between-subject variability (Zeidman et al., 2019b); all participants were retained, since PEB weights contributions by posterior certainty. A design matrix of X=[1…1]ᵀ reflected interest in group means only. To avoid evidence dilution from many parameters, separate PEB analyses were run for the A matrix and the B/C matrices (Zeidman et al., 2019b). Bayesian model reduction (BMR) with greedy search pruned weakly supported connections across the full 2^k nested model space (Friston & Penny, 2011; Pinotsis et al., 2016), and Bayesian model averaging (BMA) averaged parameters across models weighted by evidence (Friston et al., 2011). Parameters were considered reliable at posterior probability >0.95.

## 3. Results

### 3.1 Behaviour

Accuracy rates and reaction times are shown in Fig. 2. To examine the independent and combined contributions of visuo-spatial attention and semantic processing, we conducted a 2×2 repeated-measures factorial analysis with visuo-spatial attention (present/absent) and semantic (present/absent) as within-subject factors. For reaction-time analyses, we excluded errors in the ATT and COMBO conditions. We found significant main effects of visuo-spatial attention (accuracy: t₂₄ = 3.97, *p* < .001; reaction times: (t₂₄ = 9.16, *p* < .001) and semantic (accuracy: t₂₄ = 9.22, *p* < .001; reaction times: t₂₄ = 16.32, *p* < .001) factors, with no significant interaction (accuracy: t₂₄ = 0.29, *p* = .775; reaction times: t₂₄ = 0.79, p = .438). Both factors independently increased task difficulty, as reflected in lower accuracy and longer reaction times when either demand was present, with the semantic factor exerting a substantially larger impact than the attention factor on both measures. The absence of a significant interaction indicates that the costs associated with attentional and semantic processing combined additively, with no evidence of a synergistic or antagonistic effect when both demands were concurrent.

**Figure 2.**
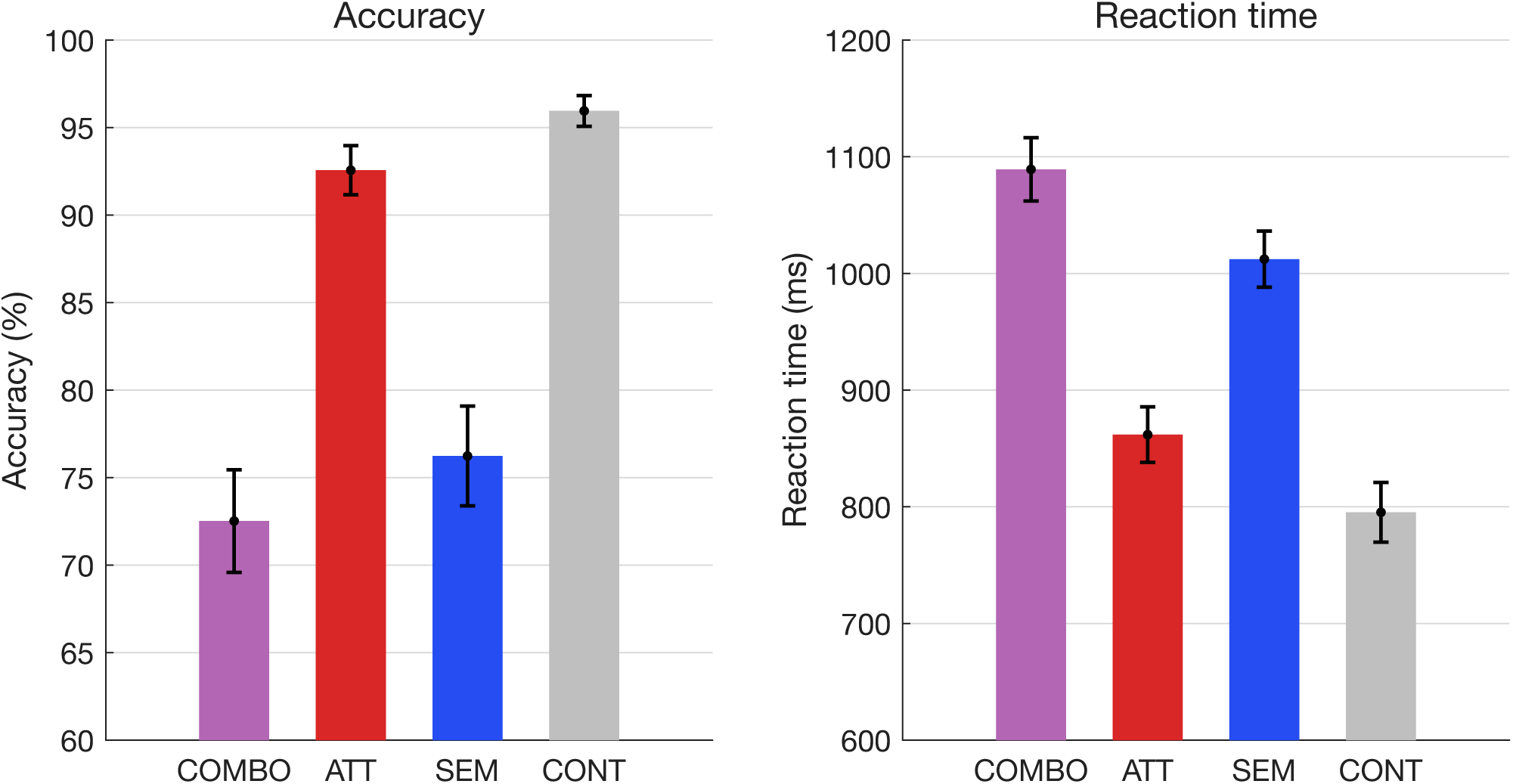
Behavioral results. Mean accuracy (%, left) and reaction time (ms, right) across the four experimental conditions (COMBO, ATT, SEM, CONT), collapsed across cue validity within ATT and COMBO conditions. Whiskers represent ±1 standard error of the mean across participants. Both accuracy and reaction time reflect increasing task difficulty from CONT through ATT to SEM and COMBO, with the semantic factor exerting a larger impact than the attention factor on both measures. For reaction times, errors were excluded in ATT and CONT conditions but retained in SEM and COMBO conditions, as words were intentionally selected to be ambiguous and force challenging semantic judgments.

### 3.2 Brain activation induced by attention

The main effect of attention ([ATT + COMBO] > [SEM + CONT]) revealed a bilateral parieto-frontal network (Fig. 3, red) comprising the frontal eye fields (FEF) and the intraparietal sulcus, peaking in the medial bank — consistent with the location of the parietal eye fields (PEF, Barash et al., 1991a, 1991b; Müri et al., 1996) — and extending posteriorly into V3A in the intraoccipital sulcus. FEF, PEF, and V3A were more strongly engaged during the attention conditions, with the highest activity in the ATT condition, followed by COMBO, CONT, and SEM. Notably, the SEM and COMBO conditions activated left hemisphere clusters more than their right hemisphere homologs, consistent with a specialization of the left hemisphere for linguistic semantic content.

**Figure 3.**
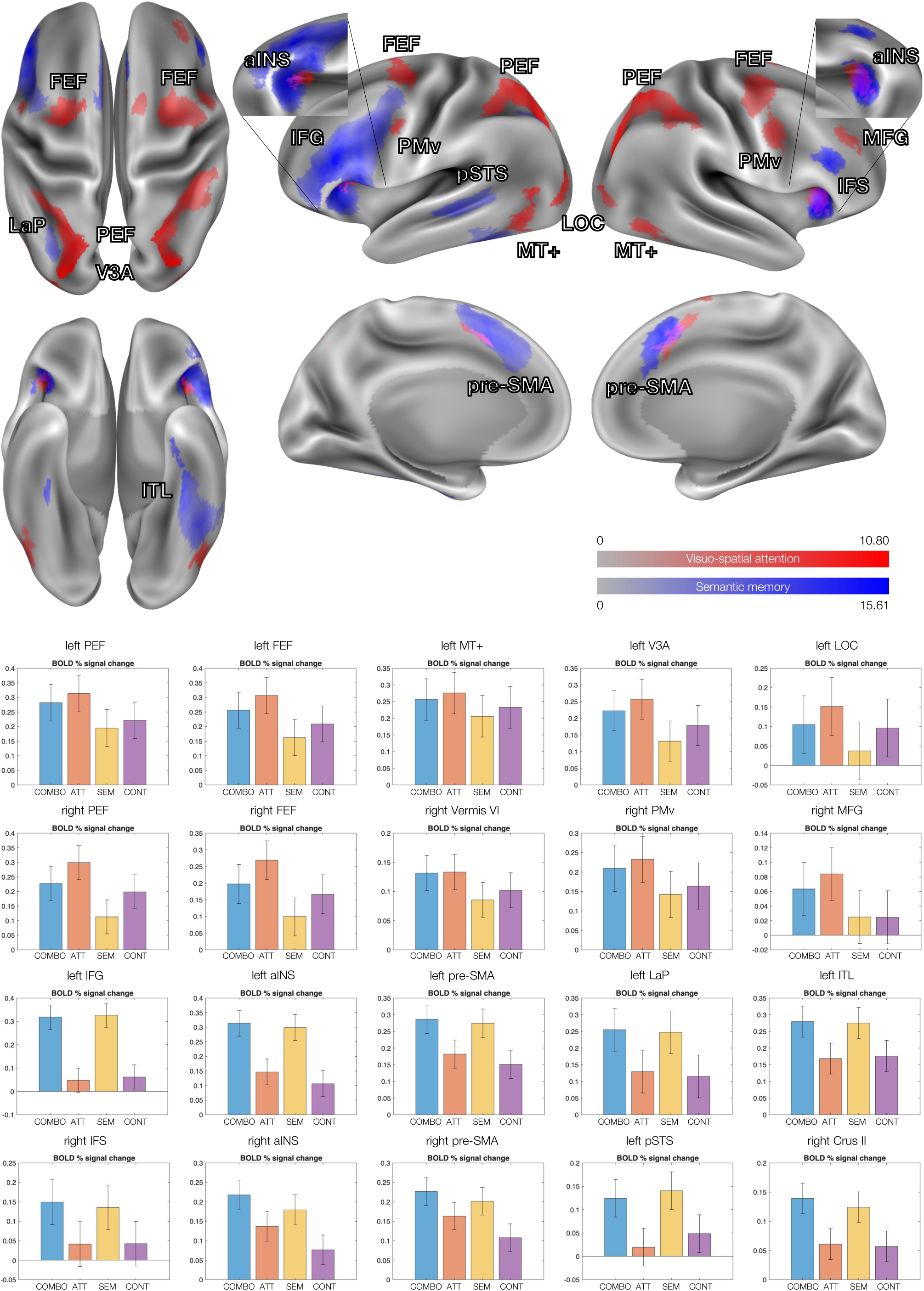
Group-level statistical parametric maps of the main effect of visuo-spatial attention ([ATT + COMBO] > [SEM + CONT], red) and the main effect of semantic processing ([SEM + COMBO] > [ATT + CONT], blue), projected onto the cortical inflated surface in fsLR space. Bar plots show mean BOLD % signal change relative to baseline across the four conditions (ATT, COMBO, SEM, CONT); whiskers represent ±1 standard error of the mean. Upper rows show regions activated by the main effect of attention (left and right PEF, FEF, MT+, V3A, LOC, Vermis VI, PMv, MFG), and lower rows show regions activated by the semantic contrast (left IFG, aINS, pre-SMA, LaP, ITL, pSTS; right IFS, aINS, pre-SMA, Crus II). Maps are thresholded at *p* < .05 corrected for multiple comparisons at the cluster level (cluster-forming threshold: *p* < .001 uncorrected at the vertex level). The attention contrast revealed bilateral activation of the frontal eye fields, parietal eye fields extending into V3A, and anterior insula. The semantic contrast revealed left-lateralized activation in the inferior frontal gyrus, the inferior temporal lobe, the posterior superior temporal sulcus, and a parietal region at the lateral bank of the intraparietal sulcus (LaP), alongside bilateral pre-SMA, the anterior insula, and the right inferior frontal sulcus. Colormaps indicate T values for each contrast; maxima of T = 10.80 for the attention contrast and T = 15.61 for the semantic contrast. Only suprathreshold vertices with average activity across conditions higher than baseline are shown. Inserts show zoomed views of activation in the anterior insula (aINS).

A similar pattern was found in the bilateral middle temporal region MT+, where activity was less differentiated across conditions. Bilateral ventral premotor cortex was also sensitive to attentional demands, particularly in the right hemisphere, with the highest activity in the ATT condition, comparable to that observed during the COMBO condition, followed by the CONT and SEM conditions.

Crucially, we found bilateral anterior insula activation, with the highest responses during the COMBO condition and the lowest during the CONT condition, with activity apparently scaling with task demands. Similarly, bilateral pre-supplementary motor area (pre-SMA) showed sustained engagement across all conditions, with higher activity during the COMBO condition, followed by SEM, ATT, and CONT, consistent with task difficulty. We also retrieved a minor activation in the right middle frontal gyrus, which showed selective recruitment during attention conditions and low responses during SEM and CONT conditions. In the cerebellum, we found activation in the left Vermis VI, with the highest similar activity during COMBO and ATT and the lowest during SEM, suggesting selectivity for attentional demands. A weaker peak was observed in the right Lobule VI, with activity scaling with task demands from COMBO to ATT to SEM to CONT.

### 3.3 Brain activation induced by semantic processing

The main effect of semantic processing ([SEM + COMBO] > [ATT + CONT]) revealed a left-lateralized network (Fig. 3, blue) comprising the left inferior frontal gyrus (IFG), which showed the strongest activation in the contrast, driven by robust recruitment during both semantic conditions against low responses during ATT and CONT. A smaller cluster in the right inferior frontal sulcus followed the same profile, at a lower magnitude.

A large cluster on the ventral surface of the left inferior temporal lobe (ITL) — including the anterior fusiform gyrus, the antero-lateral parahippocampal gyrus, and the caudal ventral inferior temporal gyrus — was strongly activated during semantic conditions. A small cluster on the right anterior fusiform gyrus showed a comparable differential pattern across conditions.

The left superior temporal sulcus was selectively recruited during semantic conditions, whereas visuo-spatial attention yielded near-baseline activity. Bilateral anterior insula (aINS) was robustly engaged, with more extensive clusters than those found in the main effect of attention. aINS effect was driven by elevated responses during semantic conditions, lower activity during ATT, and the lowest during CONT. In the left hemisphere, aINS showed comparable BOLD signal change during COMBO and SEM In contrast, in the right hemisphere, we observed significantly higher responses during COMBO than during SEM, ATT, and CONT.

Furthermore, we found activation in bilateral medial prefrontal cortex regions, including pre-SMA, encompassing larger clusters than those retrieved in the main effect of attention, extending antero-inferiorly into the anterior cingulate cortex. These pre-SMA clusters showed selectivity for semantic conditions; however, activity elicited by ATT was higher than that elicited by CONT.

Of note, bilateral caudate nucleus and left thalamus were also sensitive to semantic conditions, with very low responses in ATT and CONT. Nonetheless, a large right and a smaller left cerebellar cluster, both peaking in Crus II, showed selectivity for semantic conditions, with enhanced activity during both COMBO and SEM and low responses to ATT and CONT.

Of particular interest, we found activation in a region at the lateral bank of the intraparietal sulcus at its junction with the dorsal tip of the angular gyrus in the left hemisphere, referred to here as lateral parietal region - LaP (peak MNI coordinates: -33, -69, 45). This region showed a clear dissociation between semantic and non-semantic conditions, with comparably high responses specifically during SEM and COMBO, and low activity in both ATT and CONT. This profile indicates that LaP is selectively sensitive to the semantic factor.

LaP was close to but distinct from the activation cluster found for the main effect of visuo-spatial attention in the posterior intraparietal sulcus, which we labeled PEF (see above). A thorough examination of individual activation maps obtained with a reduced smoothing of 4 mm FWHM (Supplementary Figure 1) showed that (a) LaP was consistently activated by semantic processing in every single participant, and (b) LaP was anatomically distinct from PEF in every single participant.

### 3.4 Interaction between visuo-spatial attention and semantic processing

The interaction contrast ([COMBO + CONT] – [ATT + SEM] and its reverse) did not reveal any cluster surviving the cluster-level FDR threshold (*p* < .05 corrected), indicating that the combined condition did not produce activation patterns that were significantly greater or smaller than the sum of the two isolated conditions. This absence of a significant interaction at the neuroimaging level is consistent with the behavioral results, where no interaction was observed in either accuracy or reaction times, suggesting that the two cognitive demands were processed largely independently at the level of regional activation, with their integration occurring at the level of effective connectivity rather than through the recruitment of additional or suppressed activation regions.

### 3.5 Effective connectivity model

Cortical nodes for the connectivity model were chosen from the statistical maps of the main effects of visual attention and semantic memory, which were consistently reported in the literature (Corbetta & Shulman, 2002; Barash et al., 1991a; Bruce & Goldberg, 1985; Thompson-Schill et al., 2003; Binder et al., 2009; Noonan et al., 2013; Jedidi et al., 2021; Blackett et al., 2022; Dumitrescu et al., 2025). For visuo-spatial attention, we selected the parietal eye fields (PEF) and the frontal eye fields (FEF). For semantic memory, we selected the inferior frontal gyrus (IFG), peaking in the pars triangularis and pars opercularis, the anterior insula (aINS), a region within the inferior temporal lobe (ITL), peaking in the anterior fusiform gyrus, and a region between the dorsal tip of the angular gyrus and the lateral bank of the intraparietal sulcus (lateral parietal: LaP).

Structural connectivity considerations informed endogenous connections (A matrix): PEF was modeled as sending connections to LaP and FEF; LaP to PEF and IFG; ITL to aINS and IFG; FEF to PEF, aINS, and IFG; IFG to ITL, LaP, aINS, and FEF; and aINS to ITL, FEF, and IFG. Modulatory effects of the experimental conditions on those connections (B matrix) included a theoretically motivated subset of the endogenous connections. For the ATT condition, modulation was specified on the reciprocal connection between PEF and FEF.

For the SEM condition, modulation was specified on the reciprocal connections between LaP and IFG, between ITL and IFG, between ITL and aINS, and between IFG and aINS. For the COMBO condition, modulation was specified for all connections included in the ATT and SEM conditions, with the addition of reciprocal connections between FEF and IFG and between PEF and LaP. Direct driving inputs (C matrix) were specified as follows: PEF for the ATT condition, LaP for the SEM condition, and aINS for the COMBO condition, representing the primary entry points of task-related perturbation into the network for each experimental manipulation.

### 3.6 Effective connectivity results

Across participants, the mean variance explained by individual DCMs was 20.32%, the mean maximum connection strength was 0.479, and the mean posterior correlation among estimable parameters was 29.69%, reflecting shared posterior variance across parameter pairs. Together, these diagnostics indicated adequate parameter identifiability and model inversion. Four participants showed explained variance below the conventional 10% threshold (range: 5–8%); however, these participants were retained for group-level inference because the PEB framework takes the full posterior probability density over DCM parameters to the group level, so participants with noisier data contribute less to the group result (Zeidman et al., 2019b).

Figure 4 summarizes the effect sizes of all significant (probability > 95%; *p* < .001) effective connectivity parameters. Regarding the intrinsic effective connectivity, LaP and PEF, as well as IFG and FEF, formed bidirectional excitatory loops, with greater excitation in a ventrolateral-to-dorsomedial direction. Moderate excitatory-inhibitory coupled cross-talks mainly constituted connections across lobes. Specifically, PEF excited FEF while receiving consistent inhibition, and LaP received significant excitation from IFG and sent weak inhibitory projections back. ITL sent relevant inhibitory connections to IFG and received weak excitation. On the other hand, aINS served as a passive receiver (excited by FEF and inhibited by ITL), delivering no impactful active projections; it had bidirectional connections to IFG and unidirectional connections to both FEF and ITL, with negligible effect sizes.

**Figure 4.**
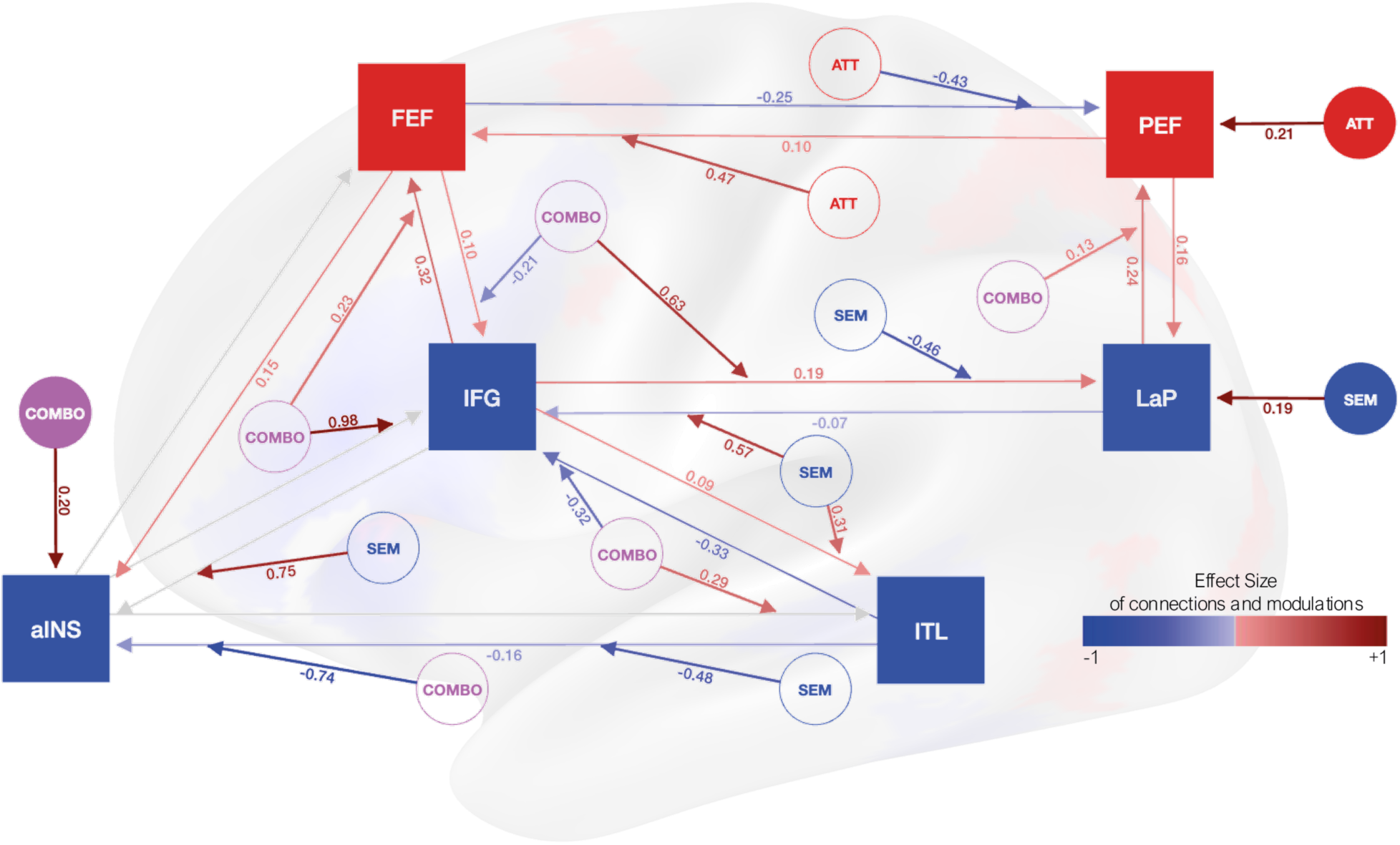
Effective connectivity results summarizing all significant DCM parameters. Filled squares represent the six ROIs — frontal eye fields (FEF), parietal eye fields (PEF), inferior frontal gyrus (IFG), anterior insula (aINS), lateral parietal region (LaP), and inferior temporal lobe (ITL) — colored according to the statistical contrast from which they were derived: red for ROIs from the main effect of visuo-spatial attention (FEF, PEF) and blue for ROIs from the main effect of semantic memory (IFG, aINS, LaP, ITL). Filled circles represent the driving inputs to the network (Matrix C), one per experimental condition. Outlined circles represent condition-specific modulatory effects on connections (Matrix B). Arrows indicate effective connections, with color ranging from blue (inhibitory) to red (excitatory) according to the color bar on the right, reflecting the sign and magnitude of the effect size; effect sizes between −0.005 and 0.005 are displayed in gray. Numbers on arrows indicate effect sizes. Conditions are color-coded as follows: red for ATT, blue for SEM, and purple for COMBO. Of note, arrows directly connecting each couple of ROIs indicate the intrinsic connectivity (Matrix A).

The driving input of each experimental condition was consistently positive on crucial regions: ATT drove PEF, the SEM condition drove LaP, and the COMBO condition drove the anterior insula. Each condition selectively modulated the effective connectivity in a peculiar pattern. ATT altered cross-talk between PEF and FEF, with strong parietofrontal forward excitation and strong frontoparietal backward inhibition. Similarly, the SEM condition modulated cross-talk between LaP and IFG, again with strong forward excitation and strong backward inhibition, thereby reversing baseline connectivity. The SEM condition also drove a critical unidirectional excitatory effect from IFG to aINS and from IFG to ITL, which, in turn, enhanced the inhibition of aINS. Such a connectivity pattern during SEM suggests a relatively independent dual-stream processing of semantic information: one relying on cross-talk between LaP and IFG, and the other on the triad of ITL, aINS, and IFG, with IFG serving as a convergence node for both pathways. The COMBO condition impacted the modulations exerted by all regions. We observed a massive propagation of excitation from aINS that excited IFG very strongly and ITL moderately. From IFG, the excitation passed to FEF and LaP, with FEF weakly inhibiting back IFG, and LaP additionally exciting PEF. From ITL, we observed inhibitory feedback on aINS, as well as inhibitory projections to IFG (see also Supplementary Figure 2).

## 4. Discussion

The present study investigated the neural mechanisms supporting visuo-spatial attention and semantic memory, both in isolation and under concurrent task demands. By employing a factorial fMRI design with effective connectivity analyses, we identified partially segregated neural systems associated with each function and characterized how these systems are reconfigured when both processes must be simultaneously engaged.

Three principal findings emerged. First, visuo-spatial attention and semantic processing recruited largely distinct cortical networks, consistent with previous literature. Indeed, regional brain activation showed additive rather than interactive effects. Second, we identified a novel region selectively and consistently recruited during semantic processing - which we term the lateral parietal region (LaP) - anatomically distinct from the parietal eye fields engaged by visuo-spatial attention. Third, dynamic causal modeling revealed substantial condition-dependent changes in effective connectivity, indicating that integration between attentional and semantic systems occurs primarily through network reconfiguration orchestrated by anterior insula, rather than by recruiting additional cortical territories.

### 4.1 Segregated parieto-frontal systems for attention and semantic processing

The main effect of visuo-spatial attention engaged a bilateral parieto-frontal network comprising parietal eye fields, frontal eye fields, motion-sensitive visual regions, premotor regions, and anterior insula, together with subcortical nuclei and cerebellar structures. This pattern closely resembles the dorsal attention network described in numerous previous studies and is consistent with the established role of frontoparietal circuits in spatial orienting and attentional control (Corbetta & Shulman, 2002; Santangelo, 2018).

The main effect of semantic memory access engaged a predominantly left-lateralized network including the inferior frontal gyrus, the anterior insula, the posterior temporal cortex, the inferior surface of the temporal lobe, and a parietal region which we labeled LaP. This activation pattern is broadly consistent with contemporary models of semantic cognition, emphasizing dependency on the specific task, stimuli and sensory modality involved (Noonan et al., 2013; Jedidi et al., 2021; Dumitrescu et al., 2025; Radman et al., 2026).

Crucially, the absence of a significant factorial interaction in both neuroimaging activation and behavioral measures indicates that the costs associated with the two demands combined additively, with no evidence of synergistic or antagonistic effects when both were required concurrently. This additive pattern is consistent with a dual-task architecture in which the two processes draw on largely dissociable resources. Of note, the semantic factor exerted a substantially larger cost than attention, consistent with semantic judgment being the dominant processing bottleneck in the combined condition.

Most importantly, at the core of each network we found a specific parieto-frontal functional coupling: a PEF–FEF loop for visuo-spatial attention and a LaP–IFG loop for semantic processing. Both networks appear to share a common functional motif characterized by parieto-frontal excitation and fronto-parietal inhibition (see also 3.3).

### 4.2 A candidate semantic parietal region at the lateral bank of the intraparietal sulcus

Among the regions associated with semantic processing, we found of great interest the area between the lateral bank of the intraparietal sulcus and the dorsal tip of the angular gyrus, referred to here as LaP. This region was consistently activated by semantic processing across participants and remained systematically distinct from the attention-related PEF region across all individual maps (see Supplementary Figure 1).

The anatomical location of LaP does not correspond straightforwardly to any commonly defined region. Its coordinates are anatomically distinct from the human parietal eye fields (the putative human homologue of the lateral intraparietal area, LIP, of the macaque), which are consistently localized on the medial bank of the IPS and are functionally characterized by their tight oculomotor coupling with FEF (Sereno et al., 2001; Schluppeck et al., 2005; Patel et al., 2010). Equally, LaP cannot be directly assimilated to the angular activations in semantic tasks which are typically reported more laterally and anteriorly (Binder et al., 2009; Seghier, 2013), nor to specific retinotopic subregions of IPS, whose visuo-spatial properties have been functionally defined (Silver et al., 2005; Swisher et al., 2007; Konen et al., 2013).

While additional work will be required to determine whether LaP represents a distinct functional area or a previously underappreciated subdivision of posterior parietal cortex, its consistent recruitment suggests that it plays a specific role in semantic processing, at least of visual material. From an anatomical standpoint, the lateral bank of the IPS is precisely where the macaque LIP is located. However, the putative human homologue of LIP has been argued to have shifted to the medial bank of the IPS during phylogenesis (Sereno et al., 2001; Schluppeck et al., 2005). This phylogenetic repositioning may reflect a competitive reorganization of cortical territory: the expansion and elaboration of the human language network — extending from inferior frontal and temporal regions into posterior parietal cortex — may have progressively recruited the lateral bank of the IPS for linguistic functions, effectively displacing the visuo-spatial attentional circuitry of LIP to the medial bank but possibly preserving oculomotor properties, which are crucial for reading. Under this account, LaP would represent a region that the language system has co-opted by recycling parietal circuitry originally serving attentional visuo-spatial functions in non-human primates (Dehaene & Cohen, 2007), with the medial bank retaining the ancestral visuo-spatial function as the human PEF.

A speculative account of LaP’s functional role can be advanced considering recent work by Hedger et al. (2025), who reported visuo-somatotopic maps in the posterior parietal cortex, in which the position of our LaP cluster falls within the cortical area corresponding to mouth and lip representation. This organization raises the possibility that LaP may serve as an integrative node linking visual and somatosensory input with the orofacial motor system. In the context of the present task, which required silent reading and semantic categorization of visually presented words, LaP may support a form of phonological simulation — the covert instantiation of articulatory or mouth-related motor programs during lexico-semantic access, consistent with embodied accounts of language comprehension (Pulvermüller, 2005; Glenberg & Gallese, 2012). This interpretation is supported by the anatomical position of LaP at the converging zone across the posterior and anterior white matter segments parallel to the arcuate fasciculus (Catani et al., 2005), the arcuate fasciculus, the inferior occipito frontal fasciculus and the superior longitudinal fasciculus III (Thiebaut de Schotten et al., 2011; Makris et al., 2005), which would place LaP at the parietal end of a circuit canonically associated with phonological processing and the dorsal language stream (Catani et al., 2005; Hickok & Poeppel, 2007).

An intriguing analogy can be drawn with the neuroscience of reach-to-grasp actions. It has been proposed that the mere perception of a graspable object activates parietal motor representations (i.e., affordances; Gibson, 1979) of the corresponding action, which are held in a state of preparatory inhibition until the goal-directed context licenses their release through frontal initiation (Jeannerod, 1994; Rizzolatti & Matelli, 2003). An analogous mechanism may operate in the domain of language: the visual presentation of a word may activate in LaP a preparatory articulatory representation — a motoric encoding of the spoken form — that is subsequently modulated by IFG, which determines whether and how the phonation affordance is expressed or suppressed. Under this framework, silent reading - as well as verbalizable thought - may be understood as inhibited speaking: a process in which the parietal instantiation of phonological-articulatory programs is continuously generated but withheld from overt execution, with LaP transforming orthographic visual input into articulatory specifications that are relayed to frontal speech and language areas for controlled deployment. This parieto-frontal loop circuitry parallels the visuomotor loop described for object-directed action, suggesting that the human brain may have recycled an ancient sensorimotor architecture to support the human capacity for language.

However, these interpretations remain speculative. The present experiment was not designed to dissociate semantic processing from covert phonological simulation, and future studies manipulating articulatory demands will be necessary to clarify the specific functional contribution of this region.

Importantly, our findings may also provide an alternative interpretation of previous TMS results from our group (Capotosto et al., 2017; 2023; 2026; Croce et al., 2018), in which stimulation targeting the angular gyrus disrupted semantic categorization performance using a similar behavioral paradigm. Given the proximity between the stimulation site and the LaP region identified here (edge-to-edge distance: ∼7 mm), those effects may have reflected interference with LaP or its underlying white matter. While this interpretation remains tentative, it highlights the importance of refining the anatomical characterization of parietal regions involved in semantic processing.

### 4.3 Network reconfiguration and the integrative role of the anterior insula

The most striking findings emerged from the effective connectivity analyses. Although behavioral and activation results suggested largely independent processing of attentional and semantic demands, connectivity analyses revealed substantial changes in network organization across conditions.

The intrinsic connectivity was characterized by significant bidirectional local excitation within both parietal (PEF-LaP) and frontal lobes (FEF-IFG) and weak excitation-inhibition cross-talks across lobes (PEF→FEF, IFG→LaP, IFG→ITL). The experimental conditions strongly modulated long-range connectivity Under isolated attentional demands, connectivity was dominated by PEF-FEF interactions, and under isolated semantic demands by LaP-IFG interactions, both characterized by forward (parieto-frontal) excitation and backward (fronto-parietal) inhibition. These findings suggest that each cognitive function relies on a dedicated parieto-frontal circuit operating as a relatively self-contained functional unit, and are compatible with the classical distinction between parietal integration of perceptual input and frontal generation of effector signals (Anderson & Buneo, 2002; Colby & Goldberg, 1999; Wolpert & Kawato, 1998; Shadmehr et al., 2010; Cisek & Kalaska, 2010).

In both loops, the parietal node can be understood supporting the transformation from sensory to motor-relevant coordinates — spatial representations in PEF, articulatory-phonological specifications in LaP — while the frontal node issues predictive signals that anticipate the sensory consequences of the planned motor output and suppress the driving parietal input. Under this account, the backward inhibition from FEF to PEF during the attention-only condition reflects a covert attentional shift signal analogous to a corollary discharge, consistent with evidence that FEF generates subthreshold motor signals even in the absence of overt eye movements (Moore & Fallah, 2001; Duhamel et al., 2005; Sommer & Wurtz, 2008), suppressing the expected inflow of information from the current parietal spatial representation. Similarly, the backward inhibition from IFG to LaP under the semantic condition may reflect a phonological corollary discharge, gating the parietal articulatory simulation once the inferior frontal speech representation has been sufficiently specified. This parieto-frontal motif finds a natural formal expression in predictive coding frameworks, in which forward connections convey bottom-up prediction errors and backward connections convey top-down predictions, with the balance between the two dynamically adjusted as a function of task demands (Friston, 2005; Rao & Ballard, 1999). To the best of our knowledge, the simultaneous observation of two such dedicated excitatory-inhibitory parieto-frontal loops within a single experimental paradigm has never been reported. It offers a uniquely controlled window into how the brain maintains functionally specialized processing streams in parallel.

The combined condition produced a markedly different pattern. Rather than simply co-activating the two circuits simultaneously, the network underwent a widespread reconfiguration involving both attentional and semantic nodes. Crucially, the anterior insula emerged as the primary driving node of this state. In the combined condition, aINS excited IFG very strongly, completely canceling the modulation observed in the semantic-only condition. Thus, aINS excitation propagated from IFG to both FEF and LaP, which also excited PEF. Simultaneously, ITL doubled its inhibition to IFG, and additionally inhibited aINS, likely regulating the overall excitatory cascade initiated by aINS.

This pattern suggests that aINS does not modulate connectivity in a diffuse or aspecific way, but rather triggers a precisely orchestrated reconfiguration of the network, in which the two task-specific loops are brought into interaction through a common hub node. At the same time, inhibitory feedback mechanisms prevent runaway excitation. The net result is a network state in which parieto-frontal coupling is enhanced within the ventral semantic loop, with fronto-parietal feedback becoming excitatory rather than inhibitory—a signature of increased cross-communication across lobes, with processed information sent back to the parietal regions.

The crucial role of the anterior insula across in the combined condition is consistent with proposals that the cingulo-opercular network — of which anterior insula is a central node — functions as a task-mode switcher or set-maintenance system operating across distinct cognitive domains (Dosenbach et al., 2007; Menon & Uddin, 2010). However, our effective connectivity results suggest a more specific and dynamic role that goes beyond generic task-set maintenance. aINS is recruited under both isolated conditions but becomes the primary driver of network reconfiguration only when the two tasks are combined. This suggests that aINS plays a crucial role specifically in dual-task performance: by propagating signals back onto both specialized networks, it may enable their concurrent recruitment, allowing otherwise competing systems to operate in parallel. From this perspective, aINS may function as a node of an independent attractor loop — possibly corresponding to the salience network — whose activation reaches a critical threshold precisely under dual-task demands, and whose dynamics can be sustained at relatively low energetic cost once established. The salience and energization functions commonly attributed to aINS (Stuss & Alexander, 2007) may therefore reflect not an intrinsic regional property, but an emergent consequence of its topological position at the ventral intersection of multiple large-scale cortical systems.

### 4.4 Functional specialization and dynamic integration

A notable aspect of the present findings is the coexistence of functional specialization and network integration: attentional and semantic processing remained largely dissociable at the regional level, yet the same networks showed substantial context-dependent reorganization in effective connectivity. This may help reconcile seemingly contradictory findings from task-based and resting-state studies. Large-scale brain networks are often described as functionally segregated systems, and several reports have emphasized competitive interactions between attention-related and default-mode regions. The present results suggest that such segregation should not be interpreted as evidence for rigid functional independence. Instead, network relationships appear highly dependent on behavioral context. When both processes were required, the same networks enter a different connectivity regime that enables coordinated processing while preserving their underlying specialization.

## 5. Conclusions

In conclusion, the present study demonstrates that visuo-spatial attention and semantic processing rely on partially segregated parieto-frontal systems that remain functionally specialized during isolated task performance but become dynamically integrated when both processes are simultaneously required. This integration is not associated with the emergence of new activation patterns but rather with a large-scale reconfiguration of effective connectivity centered on the anterior insula. These findings offer new insight into how specialized cognitive systems interact to support complex behavior, underscoring the value of studying cognition through the mechanisms that coordinate distinct functions, not only the functions in isolation.

More broadly, the present findings highlight a general challenge in cognitive neuroscience. Functional labels such as “attention” and “semantic memory” describe behavior but do not necessarily correspond to elementary neural computations. The activity observed here may prominently involve lower-level sensorimotor simulations, predictive processes, or task-specific optimization mechanisms contributing to performance. Distinguishing among these possibilities will require future studies capable of manipulating these components independently.

Several limitations should be acknowledged. First, the absence of eye-tracking data prevents a direct assessment of the contribution of covert oculomotor processes to the attentional network. Second, the DCM architecture was chosen considering current state of the literature and the findings in our specific task and therefore does not exhaust all possible network configurations. Finally, the functional interpretation of LaP remains provisional and requires direct experimental testing. Future work with novel tasks will help clarify the computations implemented by the nodes of these networks and how their reconfiguration unfolds.

## Acknowledgments

We thank Moreno Coco, Stefano Lasaponara, and Fabrizio Doricchi for their suggestions in the stimuli design.

## Data and Code Availability

All data and code will be available after acceptance of the paper, upon request from the corresponding author, subject to the need for a formal data-sharing agreement and approval from the requesting researcher’s local ethics committee.

## Author Contributions

Michelangelo Tani: Conceptualization, Data Curation, Formal Analysis, Investigation, Methodology, Software, Visualization, Writing—Original Draft, Writing—Review and Editing.

Sandeep Kaur: Investigation, Methodology.

Maria Ciociola: Writing—Original Draft.

Maria Bianca Muneghina: Investigation. Alma Cecconi: Investigation.

Valentina Sulpizio: Conceptualization, Writing—Review and Editing.

Antonello Baldassarre: Conceptualization, Writing—Review and Editing.

Paolo Capotosto: Conceptualization, Funding Acquisition, Methodology, Writing—Review and Editing.

Gaspare Galati: Conceptualization, Funding Acquisition, Methodology, Project administration, Resources, Software, Supervision, Writing—Review and Editing.

## Supplementary Material

**Supplementary Figure 1.**
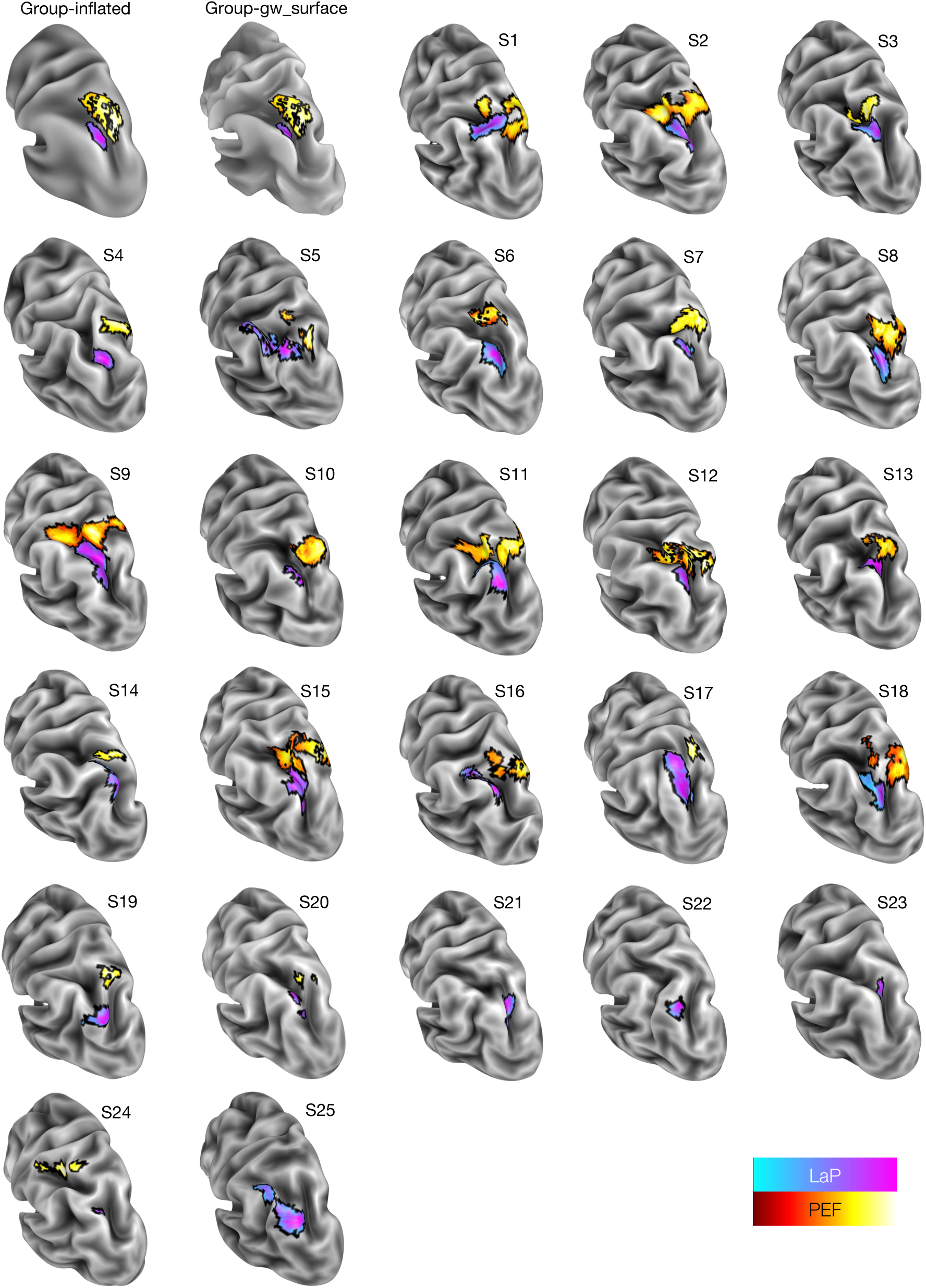
Brain mapping of LaP and PEF loci. The LaP (cool colormap, cyan to magenta), derived from the main effect of semantic memory, and PEF (hot colormap, dark red to yellow), derived from the main effect of visuo-spatial attention, are shown on the left hemisphere in a postero-lateral view, illustrating their anatomical separation at the group level and across individual participants. The first two panels show the regions on the group-level surfaces: inflated (Group-inflated) and mid-thickness (Group-gw_surface). The remaining panels (S1–S25) show the individually defined regions for each of the 25 participants on their native cortical mid-thickness surface. LaP is consistently located between the dorsal tip of the angular gyrus and the lateral bank of the posterior intraparietal sulcus. In contrast, PEF is located more superiorly on the medial bank of the intraparietal sulcus, confirming their dissociation across participants.

**Supplementary Figure 2.**
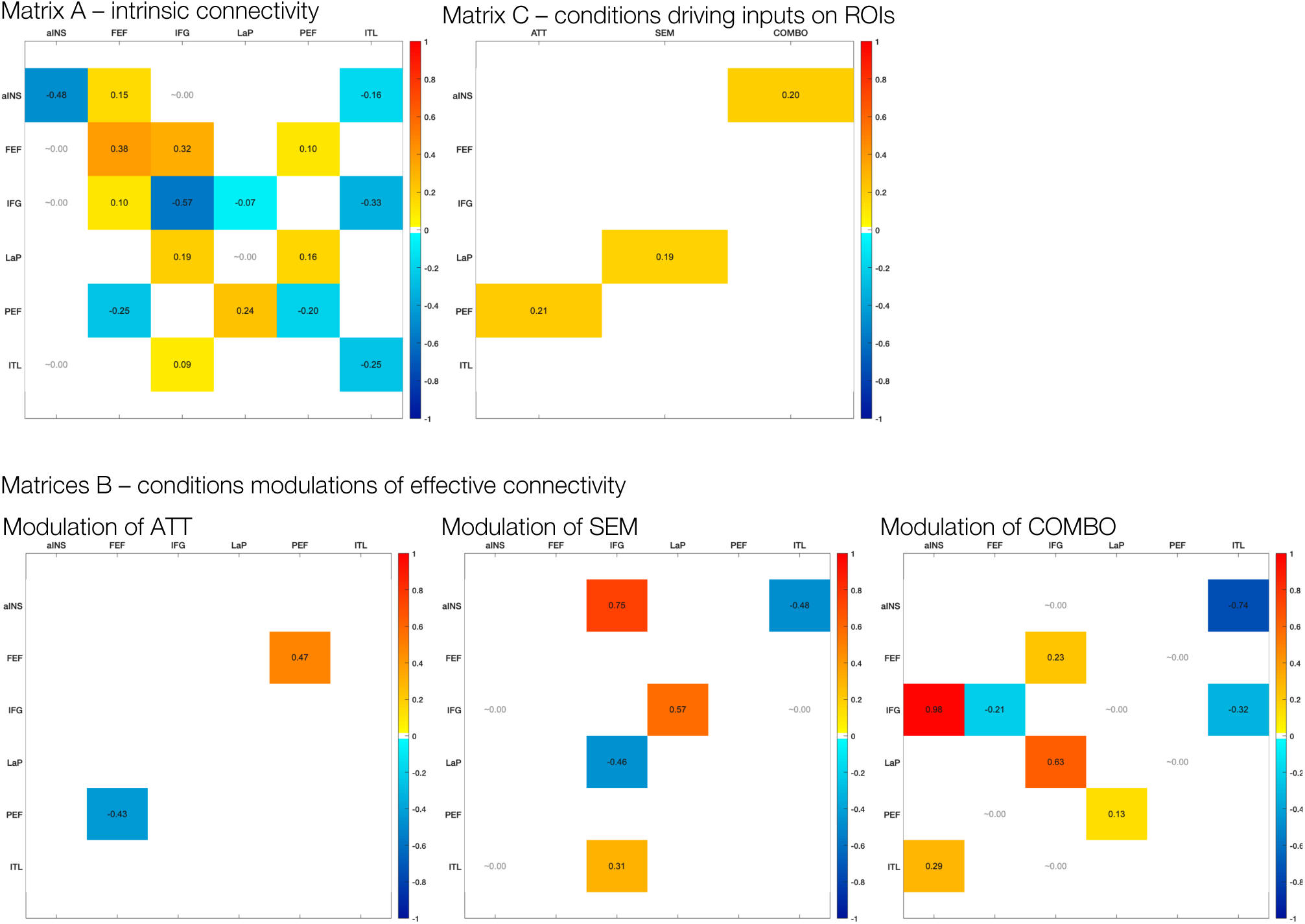
DCM parameter matrices. Matrix A shows intrinsic effective connectivity across all conditions; Matrix C shows the driving input of each experimental condition onto the network entry points; Matrices B show condition-specific modulations of effective connectivity for the attention (ATT), semantic (SEM), and combined (COMBO) conditions separately. In all heatmaps, only parameters with a posterior probability above 95% are displayed; warm colors indicate excitatory effects, and cool colors indicate inhibitory effects, with color intensity reflecting effect size.

**Supplementary Table 1.** Cluster and peak statistics for the main effect of visuo-spatial attention ([ATT + COMBO] > [SEM + CONT]). For each cluster, the table reports hemisphere, cluster size (number of vertices), cluster-level uncorrected p value, FWE-corrected p value, and topological FDR-corrected p value, followed by the MNI coordinates (x, y, z), T statistic, Z score, and uncorrected, FWE-corrected, FDR-corrected, and topological FDR-corrected p values for each local maximum. Clusters are ordered by size in descending order.

| Structure | Size | ClusterTopoFdrP | x | y | z | T |
| --- | --- | --- | --- | --- | --- | --- |
| Right V3A | 1032 | 5.65977E-75 | 33 | -71 | 26 | 10.7992566 |
| Right PEF |  |  | 25 | -61 | 55 | 8.56802516 |
| Right aIPS |  |  | 43 | -37 | 43 | 5.8395657 |
| Left V3A | 875 | 1.01297E-63 | -29 | -82 | 20 | 8.48309992 |
| Left PEF |  |  | -21 | -65 | 51 | 7.0245962 |
| Left aIPS |  |  | -36 | -43 | 44 | 4.25516161 |
| Left LOC | 114 | 1.85428E-08 | -31 | -87 | 17 | 6.83628725 |
| Right MT+ | 125 | 3.51416E-09 | 47 | -63 | -6 | 6.24715986 |
| Right LOC | 61 | 0.000101341 | 33 | -85 | 11 | 6.04768819 |
| Left superior LOC | 188 | 1.21805E-13 | -44 | -69 | 12 | 5.96447863 |
| Left inferior LOC |  |  | -41 | -73 | -12 | 4.22068247 |
| Right PMv | 156 | 2.06845E-11 | 47 | 3 | 35 | 5.78173704 |
| Right FEF | 378 | 2.53721E-27 | 29 | 0 | 54 | 5.77835631 |
| Left Cerebellum Vermis VI | 228 | 3.72638E-05 | -4 | -74 | -24 | 5.55542515 |
| Right Cerebellum Lobule VI | 103 | 0.00232567 | 10 | -78 | -18 | 5.24461283 |
| Right MFG | 90 | 9.15828E-07 | 33 | 42 | 25 | 4.96229747 |
| Right aINS | 173 | 1.32613E-12 | 32 | 27 | 3 | 4.94459846 |
| Left FEF | 219 | 7.64744E-16 | -25 | -8 | 51 | 4.92996402 |
| Left aINS | 85 | 1.9842E-06 | -31 | 21 | 11 | 4.92251511 |
| Right PMd | 66 | 4.63293E-05 | 10 | 3 | 70 | 4.74619888 |
| Right pre-SMA | 115 | 1.72177E-08 | 7 | 9 | 54 | 4.58983022 |
| Left PMv | 36 | 0.006595323 | -46 | 0 | 36 | 4.25922216 |
| Left pre-SMA | 33 | 0.010355861 | -6 | 12 | 45 | 4.09045902 |

**Supplementary Table 2.** Cluster and peak statistics for the main effect of semantic processing ([SEM + COMBO] > [ATT + CONT]). For each cluster, the table reports hemisphere, cluster size (number of vertices), cluster-level uncorrected p value, FWE-corrected p value, and topological FDR-corrected p value, followed by the MNI coordinates (x, y, z), T statistic, Z score, and uncorrected, FWE-corrected, FDR-corrected, and topological FDR-corrected p values for each local maximum. Clusters are ordered by size in descending order.

| Structure | Size | ClusterTopoFdrP | x | y | z | T |
| --- | --- | --- | --- | --- | --- | --- |
| Left IFG | 1735 | 8.4039E-131 | -42 | 29 | 18 | 15.6127382 |
| Left aINS |  |  | -30 | 28 | 5 | 11.6658845 |
| Right Cerebellum Crus II | 597 | 1.35642E-08 | 10 | -84 | -32 | 11.6642294 |
| Right Cerebellum Crus I |  |  | 12 | -74 | -28 | 7.7106884 |
| Left ITL anterior fusiform gyrus | 415 | 2.69094E-31 | -37 | -30 | -24 | 9.32664928 |
| Left ITL ITG |  |  | -53 | -56 | -13 | 4.26158478 |
| Right aINS | 276 | 6.24358E-21 | 37 | 20 | -2 | 9.11773071 |
| Left pre-SMA | 473 | 1.43278E-35 | -6 | 17 | 46 | 8.95976588 |
| Right Cerebellum Crus II | 134 | 0.001189772 | 34 | -68 | -44 | 8.24947508 |
| Right pre-SMA | 233 | 8.54225E-18 | 6 | 25 | 44 | 8.043452 |
| Right IFS | 122 | 1.37673E-09 | 45 | 30 | 18 | 7.82135319 |
| Left parahippocampal gyrus | 39 | 0.00235146 | -30 | -5 | -38 | 7.69618004 |
| Left Caudate | 246 | 4.33171E-05 | -14 | -2 | 16 | 7.53553876 |
| Left Thalamus | 230 | 5.59453E-05 | -8 | -14 | 6 | 7.12288064 |
| Left pSTS | 338 | 1.48798E-25 | -50 | -31 | -2 | 6.75936558 |
| Left LaP | 142 | 4.95016E-11 | -33 | -69 | 45 | 6.50204705 |
| Left Cerebellum Crus II | 81 | 0.009198158 | -6 | -82 | -32 | 6.02668512 |
| Right Caudate | 150 | 0.000781596 | 14 | 12 | 6 | 5.35010796 |
| Right ITL anterior fusiform gyrus | 25 | 0.025412941 | 35 | -30 | -25 | 4.91598083 |

